# Foraging analysis of Endangered Greater Adsjutant Stork *Leptotilus dubios* Gemlin in certain habitat of Assam, India

**DOI:** 10.1101/2020.05.31.125328

**Authors:** Purnima Devi Barman, D. K. Sharma

## Abstract

The endangered Greater Adjutant Stok *Leptotilus dubius* Gemlin confined only in some pockets in Assam and Bihar in India and in certain areas of Laos and Cambodia has poorly understood in its foraging activities in its natural habitats. Attempt has been made to address the forage pattern in its natural habitat wetlands in the Kamup district and in a city garbage dump in Guwahati in Assam, India. The various forage methods like walking, visual tactile etc and their relation with the body metrices like beak length, tibia tarsus length and its mass were assessed in terms of foraging habits. Forage actvities were evaluated during the breeding (May to September) and non breeding (October to April) time of the study period of 2012-2017 at different water level of the wetland(s). The foraging range of this stork assessed at an confinement of aerial distance of 15 km from the nesting sites appears as resident non migratory birds by habits. Types of diet composition obtained from the regurgitated food at the ground of the nesting sites were mostly the fish group channa and cyprinids. Profitability index was determined at the captive stork showed in favour of these two groups within the size group of 5 to 15 cm. Larger food item showed lesser profitability index. PCA analysis showed negative foraging correlation with the Prey size greater than 8cm and 15 cm, while the captivity study was conducted Thus the present findings on the foraging assessment of Greater Adjutant might be the baseline information for conservation action plan.

## INTRODUCTION

The endangered Greater adjutant stork *Leptoptilos dubius* (Luthin, 1987; IUCN, 2018) appears as global stronghold species (Jetz *et al*.,2014) with 800 birds in the Kamrup district of the Brahamaputra valley of Assam (Singha *et al*., 2003, Goswami and Patar, 2007, Barman *et al*., 2015, Birdlife International,2018) and in Bihar in India (Saikia and Bhattacharjee, 1989a; Rahmani, 1989; Rahmani *et al*., 1990; Hancock *et al*., 1992; Choudhury *et al*., 2004; Mishra and Mandal, 2009) and few in South East Asian countries (Luthin, 1987; Campbell *et al*., 2006; Clements *et al*., 2007). The Greater Adjutant stork (GAS), locally called as *Hargila* (the bone swallower), *Hodong, Dhodong, Jomtokola and Bamuni bortokola* etc. (Singha, 1998) and named as “adjutant’’ because it walks with deliberate measured gait of a military Adjutant (Singha, 1998).Being carnivorous, it occupies the top position in the food chains, plays significant role in wetland ecosystem (Singha, 1998), but with little or poor information (Saikia and Bhattacharjee, 1989a,1989b, Rahmani *et al*., 1990), despite they use to forage in 50 different wetlands in the Kamrup district of Assam. Furthermore, the district supports 300 to 450 birds together as foraging habitat in a garbage dumping ground but without details on foraging ecology, habitat utilization and breeding behaviour of the species GAS attracts attention for conservation due to its decreasing population trend (Luthin, 1987, Singha *et a*l., 2003a).

Foraging ecology of birds resolves around making use of diverse foraging opportunities in space and time (Kushlan,1981). Feeding behaviour is the main factor that affects the foraging efficiency in wading birds (Whitfield and Blaber, 1979). Foraging ecology of stork (Ogden *et al*.,1976) is governed by seasonality of prey ((Kahl 1964, Coulter, 1987), prey resources, habitat characteristics(Maheswaran and Rahmani, 2002), seasonal rainfall (Odum *et al*., 1995) foraging organs (Kahl 1964, Kushlan 1981, Coulter and Bryan 1993, Urfi, 2011). Food preferences and feeding techniques are related to the individual’s morphological features, particularly the size and shape of the bill used for seizing food, lengths of neck and leg, which determine its foraging efficiency (Hancock and Kushlan, 1984) and reproductive success (Bryan *et al*., 1995) and the colonial nesting bird, owing to their long breeding season, requires huge amount of food to feed their nestlings. The Greater Adjutant by habit mostly forages in wetlands (Saikia and Bhattachajee, 1989a; Newton, 1998, Singha *et al*., 2003, Barman *et al*., 2015), constructs their nesting colonies to the close proximity of food and its variation affects both fecundity and reproductive success including the number and quality of young (Martin,1987; Newton,1998). Thus an attempt has been made to assess the foraging behaviour of Greater Adjutant Stork within their radial distance of habitat locations

## Materials and Methods

The present work was conducted in 6 wetlands and in a garbage dump in the Kamrup District of Assam, India situated between 25°43/ and 26°51 N Latitudes and 90°12/ and 90°36/ E Longitudes (Table 1). Part of the study was also performed in captivity in the Assam state Zoo. Climate wise the cold season starts from November to February followed by pre-monsoon from March to May, monsoon from June to August and post monsoon from September to October(Borthakur,1986). The average annual rainfall in the district is recorded at 2135.7 mm ranging between 1500 mm to 2600 mm in the Brahmaputra valley and the mean daily maximum and minimum temperature stands at about 25.2°C and 10°C during winter. The investigation period (2012-2017) has been divided into a) breeding season from April to September and b) non breeding season from October to March in a given year.

**Table 1:**
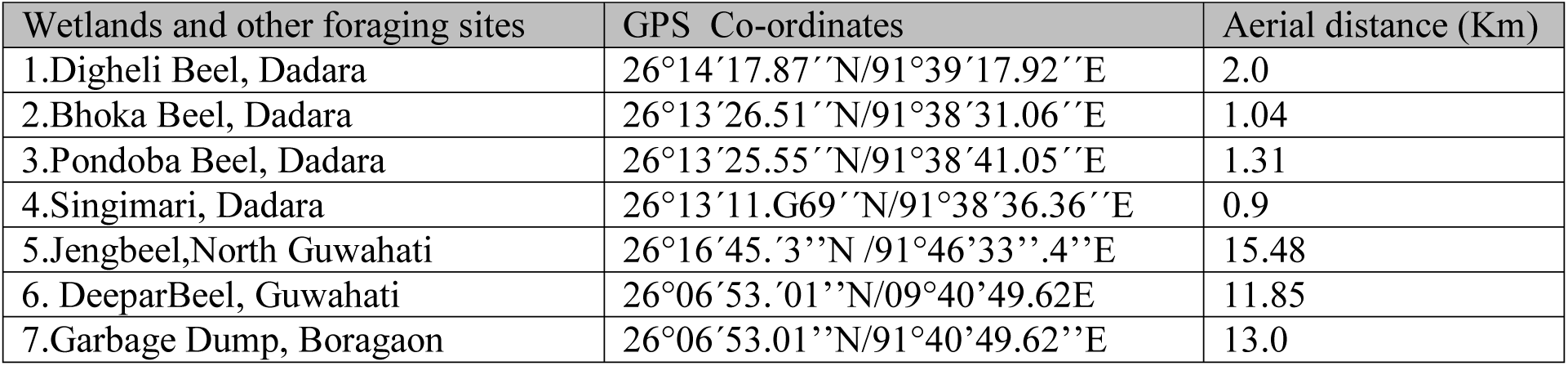
GPS coordinates of foraging wetlands and other foraging site of Greater Adjutant Stork.

Selection of Foraging sites: Survey was conducted from 10th January 2012 to 10^th^ January 2013 to locate the major foraging grounds in the Kamrup District. The foragers were counted thrice in a week and the mean number of birds per week was estimated. Later the wetlands which had more than 50 birds (mean number), were selected as primary foraging sites. Aerial distance between foraging ground and nesting location from Dadara, Pachariya, Singimari village and major wetlands used were estimated by taking the help of GIS laboratory of Aaranyak (Google Earth Pro, Version 7.0, Google INC 2012). The location of each study point was stored in Garmin GPS and a GIS map was prepared showing the aerial distance of foraging grounds and breeding ground (Table-1)

Body morphometry: Morphometry was recorded both in live and dead birds separately and opportunistically. Records on live birds were done in captive and rescued birds with the support from Veterinarian and expert from the Assam State Zoo. Since the live birds sample were less, morphometry (Urfi,2011) was separately done in died birds, that too opportunistically to get more sample size of 6 adult and 14 sub adult dead as well as 6 adult and 6 sub adult live birds on the following parameters: Beak length/bill length and opening length: Beak length or bill length is estimated as distance between tip of the upper mandible and the other at base of the skull.

Tibia length: Tibia length is estimated as the distance from joint of tibia-tarsus joint and foot.

Tarsus length: Estimated as distance from tibia tarsus joint and foot.

Body length: Estimated as distance between tibia tarsus joint and foot.

Body Depth: Estimated as the distance from the highest point on the back to the belly.

Recording of Water depth in wetlands : For each individual, water depth at the foraging site was estimated by noting the point up to which its legs were submerged (Maheswaran and Rahmani, 2002). Depth at wetlands were recorded by measuring tap and calibrated scale at every week interval. Based on the leg size and direct measurement, the water depth were divided into: 1 to 10 cm (S_1_), 1 to 30 cm (S_2_), 1 to 40cm (S_3_), 1-50cm S_4_) and 1 to 70cm and above(S_5_). Data were obtained twice in a week at different wetlands from 5:00 a.m to 6:00 p.m. The day was divided into four blocks 5:00 am to 9:00pm, 9 am to 12:00, 12:00 to 3:00pm, 3:00 to 6:00pm. 5 min was considered as one bout.

Foraging behaviour and strategies: Observation on foraging behavior was recorded after Gonzalez, (1997) using 10×50 binoculars or a 20X spotting scope from January 2012 to January 2017. The year was divided into two periods: a breeding season (April to September) and a non breeding season (May to October). Foraging behaviors were recorded as probing, pecking, groping and head swinging (Kushlan,1977, Ishtiaq *et al*., 2004).Probing is the insertion of the slightly open mandible deep into the sediment, under plant roots or rocks. Pecking is the picking up of sighted objects without inserting the bill into the substrate. Groping is holding a widely gaping bill in the water while moving the tip along the bottom. Head Swinging is moving the partially submerged and gaping bill from side to side in the water. Foraging was observed for a mean period of 12 days per month. Prey handling methods (Khuslan, 1977; Maheswaran and Rahmani, 2002) and foraging techniques (Dorfman *et al*.,2001) were adopted. Focal animal sampling was used to record the behaviors of Greater Adjutant Stork (Altman, 1974). The following parameters were considered for the study

a. Foraging bout: Starting and ending time of each foraging bout
b. Number of pecks/probes was counted with a hand tally counter.
c. Number of fish caught: Number of fishes caught per 5 min bout was later converted into fish caught per minute
d. Prey size: Prey size was measured against the size of the bill into classes as one fourth, half of the bill or full size of the bill (Maheswaran and Rahmani, 2002)
e. Prey handling time: Refers the time required from time of capturing the prey until it swallowed.
f. Water depth: The depth of water in various wetlands in cm, where the birds foraged.
g. The mode of food capture: The visual and tactile method of foraging.
h. Peck rate: Refers to the number of attempts made by the bird. It is defined as number of times the stork jabbed the water with its bill in search of prey.
i. Feeding rate(FR): Feeding rate is the total number of feeding attempts (pecks and probes) made per minute per bird.
j. Feeding success rate: Number of fishes caught per minute divided by number of foraging attempts per minute+ number of probe.

Prey handling time (PHT) : Prey Handling Time (PHT) was recorded in both wild and captivity by observing the birds using 50 × 10 binocular from a distance of 50m in field (approximately) keeping time records in a stopwatch. Each observation of handling a fish by a stork was considered as one bout. The sizes were based on comparison on the half the size of beak, one fourth of the beak and full size of the beak (Maheswaran and Rahmani 2002). Since the sizes were based on assumption and the same was also carried in captivity in the Assam State Zoo on 50 samples of known fishes for profitability index.

Profitability Index : The cost of eating each prey item was estimated by its handling time (prey handling time in s) and the benefit was estimated as the dry weight of the item as depicted in the formula (Urfi, 2011). Profitability index= Dry weight of prey item (g)/ Prey handling time (s)

## Results

Morphometry : Sexual dimorphism could not be recorded for Greater Adjutant stork (though the lager and taller is considered as the male during nesting). Body morphometry of live birds was recorded in captivity depicted in Table 2, represents that tibia and tarsus of adult bird significantly different (P<0.005) from that of the sub adult. Records of forages at various depth level and prey catches are presented in the Table-3.

**Table 2:** Body morphometry (in cm) of Greater Adjutant Stork (CV=Coefficient of Variation); ± = SD. Figures in the parenthesis indicate range variation. * t test (p<0.05)

| Parameters | Dead |  |  |  | Live |  |  |  |
| --- | --- | --- | --- | --- | --- | --- | --- | --- |
|  | Sub Adult (14) | CV | Adult (3) | CV | Sub Adult (6) | CV | Adult (6) | CV |
| Body Depth | 30.96±1.31<br>(25-38) | 4.2 | 33.93±2.28<br>(29-40) | 6.71 | 33.96±1.43<br>(29-38) | 4.2 | 33.00±0.0<br>(33 to 33) | 0 |
| Bill/beak length | 29.20±1.75<br>(28 to 30) | 5.9 | 36.23±1.20<br>(33-37) | 3.97 | 30.10±0.75<br>(28 to 30) | 2.49 | 38.22±1.31<br>(37 to 38) | 3.42 |
| Beak opening | 25.46±1.62<br>(20-30) | 6.3 | 29.83±1.84<br>(25-38) | 6.16 | NA | NA | NA | NA |
| Head length | 9.15±0.88<br>(7-10) | 9.61 | 10.54±4.76<br>(8-11) | 7.2 | 10.15±0.22<br>(7-10) | 2.49 | 12.00±0.0<br>(12-12)) | 0 |
| Head breadth | 7.21±0.57<br>(5-9) | 7.9 | 8.16±2.93<br>(6.5-10) | 2.5 | 78.11±0.11<br>(5-10) | 0.14 | 8.54±0.32<br>(8-9) | 3.74 |
| Body length (head to body end) | 57.34±1.62<br>(50-65) | 2.8 | 62.57±1.93<br>(63-75) | 4.68 | 64.44±0.2<br>(50-65) | 0.31 | 86.45±0.40<br>(65- 88) | 0.46 |
| Body length (Head to tail) | 95.64±2.81<br>(85-100) | 2.9 | 105.4±4.28<br>(90-120) | 4.06 | 98.44±2.12<br>(83-100) | 2.14 | 105.4±3.09<br>(90-110) | 2.93 |
| Wing span | 295.24±3.31<br>(260-300) | 1.12 | 310.51±3.11<br>(290-340) | 1.00 | 299.92±3.02<br>(280-300) | 1.0 | 310.11±3.11<br>(290-340) | 1.0 |
| Tarsus | 32.24±2.93<br>(26-38) | 9.63 | 46.94±3.12*<br>(40-50) | 9.47 | 34.47±2.22<br>(30-38) | 6.44 | 46.54±3.87*<br>(42-51) | 8.31 |
| Tibia | 37.33.1±2.45<br>(35-43) | 5.9 | 43.54±2.39*<br>(39-45) | 7.05 | 36.54±2.45<br>(32-43) | 6.7 | 43.54±2.39*<br>(39-45) | 5.48 |
| Full leg | 68.20±1.23<br>(65-99) | 1.75 | 85.10±1.98<br>(82-89) | 2.75 | 68.99±2.03<br>(61-87) | 2.94 | 86.02±1.33<br>(82-89) | 1.54 |
| Body Weight (g) | 7.24±1.11<br>(6-9) | 8.76 | 08.56±0.21<br>(7-9) | 2.19 | 7.77±1.01<br>(5- 8.5) | 9.76 | 8.01±0.22<br>(7-9) | 1.83 |

**Table 3:** Numbers of birds at different water depth(s) in different months (as%) both in breeding and non breeding season during the period of 2012 to 2017

| Month | S1(18%) | S2(21%) | S3(26%) | S4(21%) | S5(14%) |
| --- | --- | --- | --- | --- | --- |
| May | 22.55±2.6<br>(10 – 30) | 40.90±3.8<br>(14 – 54) | 35.70±3.3<br>(14 – 49) | 33.80±1.8<br>(29 – 40) | 9.30±1.01*<br>(7 – 15) |
| June | 26.25±3.6<br>(19 – 49) | 41.30±2.8<br>(27 – 58) | 48.20±4.6<br>(30 – 60) | 31.34±2.0<br>(26 – 48) | 14.17±1.89*<br>(9 – 19) |
| July | 40.34±3.1<br>(30 – 45) | 48.44±3.6<br>(20 – 50) | 57.11±5.45<br>(40 – 30) | 49.32±4.04<br>(30 – 56) | 11.40±4.50<br>(9 – 15) |
| Aug | 61.64±4.7<br>(30 – 70) | 62.30±5.0<br>(30 – 86) | 57.0±4.60<br>(40 – 75) | 34.94±2.6<br>(28 – 63) | 19.40±5.32**<br>(11 – 42) |
| Sep | 41.45±2.6<br>(41 – 75) | 41.25±3.1<br>(35 – 70) | 58.40±4.1<br>(1 – 96) | 34.87±3.7<br>(28 – 75) | 30.45±4.25<br>(31 – 90) |
| Oct | 65.50±6.1<br>(40 – 100) | 68.24±5.7<br>(48 – 105) | 59.34±4.1<br>(45 – 100) | 67.85±4.3<br>(48 – 112) | 30.34±5.16<br>(18 – 40) |
| Nov | 46.48±3.4<br>(40 – 90) | 65.30±1.0<br>(50 – 105) | 56.24±1.9<br>(50 – 86) | 59.15±3.3<br>(48 – 95) | 33.68±5.42<br>(25 – 55) |
| Dec | 48.42±3.2<br>(42 – 83) | 50.24±3.6<br>(41 – 90) | 73.10±5.0<br>(51 – 105) | 49.0±2.30<br>(42 – 86) | 31.26±4.2<br>(25 – 55) |
| Jan | 24.31±1.1<br>(19-28) | 36.33±1.9<br>(21-40) | 33.45±2.2<br>(18-60) | 28.16±1.8<br>(20-30) | 24.15±1.9<br>(18-31) |
| Feb | 46.47±1.9<br>(16-80) | 58.34±2.8<br>(26-70) | 80.11±1.7<br>(22-106) | 61.22±1.7<br>(23-77) | 21.34±2.1<br>(10-30) |
| March | 18.45±2.2<br>(17-26) | 29.31±1.9<br>(20-38) | 25.16±1.6<br>(16-30) | 28.34±1.9<br>(18-36) | 20.24±1.62<br>(22-34) |
| Apr | 22.38±1.9<br>(17-26) | 28.47±2.21<br>(21-66) | 65.21±2.16<br>(28-100) | 58.31±1.72<br>(24-66) | 26.15±2.34<br>(18-32) |

(Observation days, n = 10/month); ± SE, Figures within Parenthesis indicate range variation. Depth at various wetlands, S_1_=1 – 10cm :S_2_=1– 30 cm :S_3_=1 – 40 cm :S_4_=1 – 50 cm :S_5_=1 to 75cm and above

Prey handling time (PHT): Different time spent by GAS in prey items of different sizes was observed as the PHT increases with the size of prey (measured by ocular method and by comparing the beak size). The PHT varied from half minute for small prey items up to maximum eighteen minutes for larger prey like snakes (Table 4).

**Table 4 :**
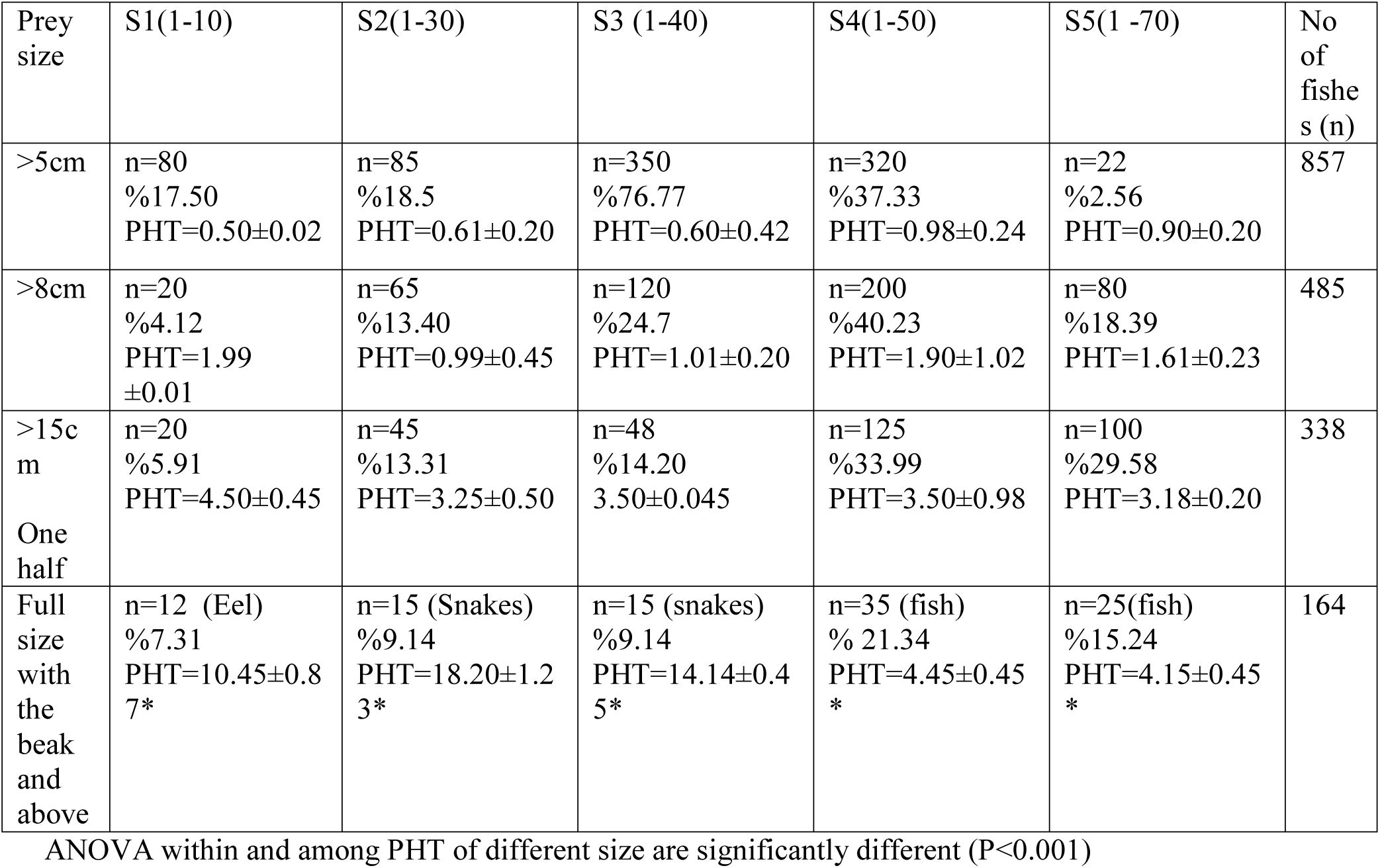
Prey handling time (PHT in % min) of Greater adjutant stork for different sizes of prey(s) and numbers of catch(%)at different water depth (S1, S2, S3, S4, S5 in cm) in wetlands during the study period. (Mean ±SE for PHT)

The PHT time increased according to size at various depth level to maximum of 18.20 ± 1.23% for snake(n=15). The one way ANOVA within and among prey handling time for different sizes of fishes were significantly different (P<0.001). Pearson correlation coefficient between PHT and other variables like, water depth (r=0.749, R^2^ =0,561) fish size (r=0.9391, R^2^ =0.8819) water surface temperature (r= 0.1982, R^2^ =0.0393) air temperature (r= −0.3231, R^2^ =0.1044), group size(r= −0.3303, R^2^ =0.1091) and other wading birds nearby (r= 0.0112, R^2^ =0.0001) The PHT is positively correlated (P<0.01) with fish size and water depth. The PHT in captivity for fish (n=50 of 6 cm, n=50 of 10 cm, n=20 of 15 cm) was 1.2 ±.04, 2.12±.09 % 3.11±.07 against Eel (n=20 of 35 cm) for 7.23±.05% minutes

Diet Composition: The diet composition of GAS in wetlands comprised of varieties fishes, birds, amphibian, reptiles and sometimes bird. Most of the prey items were obtained as regurgitated food below the nesting tree (Table 5). 426 prey items could be collected during the study period and calculated for group Channa 20.51%, Cyprinus 16.86%, Notopteros 5.28% cat fish (*H*.*fossilis, C. magur*) 22.27%, Eel 2.24%, frogs 3.68% and others at 15.84 % including Crab, Snake, Gallus Snails etc. Further, the diet spectrum comprised of animal viscera, remains of fish from the fish market, carcasses, chicken feathers, remains of butcher houses, cooked meat wastage from the restaurant etc were recorded from the gut analysis of dead bird.

**Table 5:**
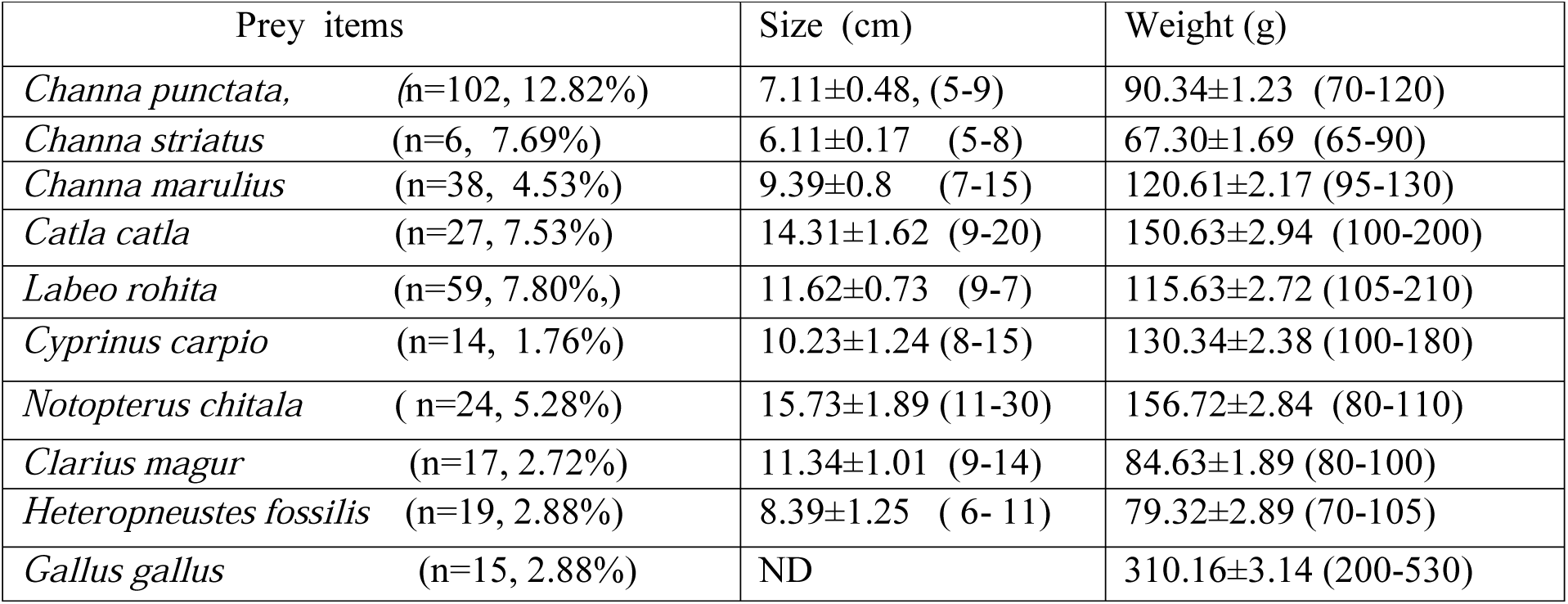
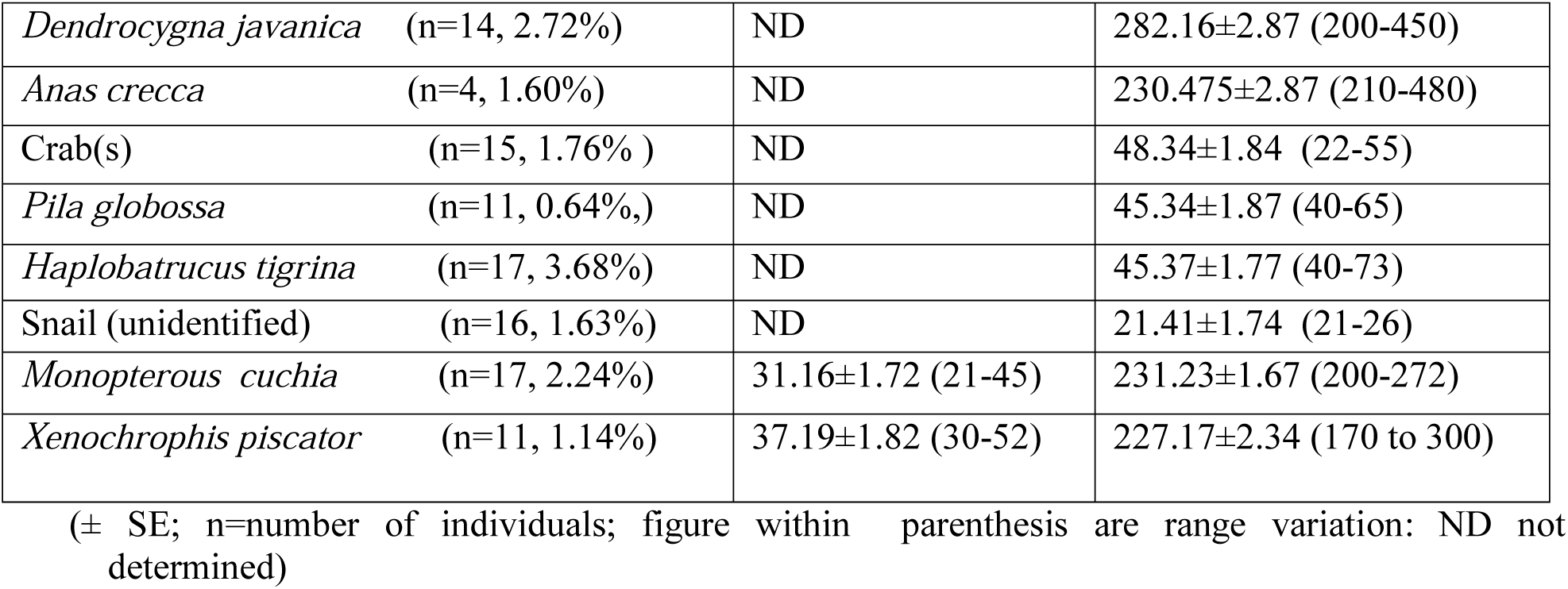
Size (cm) and weight (g) of different regurgitated prey component as % in diet spectrum of collected under the nesting site(s) of Greater Adjutant Stork during the breeding period (2012-2017

Greater Adjutant Storks arrived at garbage dump at around 4:30 am and start foraging from 5:30 am but was found to be more active from around 6 am In the garbage dump, storks became very much adapted to the arrival of garbage trucks. As soon as the garbage truck arrive, they congregate in the garbage and their feeding frequency increases

Profitability index:The fishes considered, were *Puntius puntius, Labeo rohita, Channa striatus, Channa marulius, Channa punctata, Monopterous cuchia, Heteropneustis fossilis*. The profitability index was carried out for adult, chicks and sub adult in captivity, Assam state Zoo (Table 6)

**Table 6:** Profitability index of certain fishes foraged by Greater Adjutant Stork in captivity.

|  | Prey handling time(seconds) and profitability index (P) |  |  |
| --- | --- | --- | --- |
| Prey item | Adult | Chick (3 to 4 months) | Sub adult |
| *Puntius puntius*<br>(n=18, 10.12) | 1.53±0.04<br>(P=6.5) | 1.62±0.01<br>(p=6.17) | 1.3±0.02<br>P=7.6 |
| *Labeo rohita*<br>(n= 16, 20.03) | 1.72±0.03<br>(P=11.65) | 1.62±0.03<br>P=12.34 | 1.62±0.02<br>P=11.95 |
| *Channa striatus*<br>(n=16, 14.64) | 1.05±0.0.2<br>(p=13.95) | 2.06±0.05<br>P=14.56 | 2.51±0.0.3<br>P=11.95 |
| *Channa marulius*<br>(n=26 25.15) | 1.04±0.0.2<br>(P=24.19) | 2.92±0.0.3<br>P=10.24 | 2.14±0.01<br>P=12.90 |
| *Channa punctata*<br>(n=26 , 26.52) | 1.41±0.0.3<br>(P=18.81) | 1.39±0.02<br>P=20.41 | 1.17±0.04<br>P=4.84 |
| *Monopteros cuchia*<br>(n=26, 17.56) | 5.15±0.05<br>(P=3.41) | 4.27±0.04<br>P=3.78 | 5.16±0.04<br>P=4.84 |
| *Heteropneustis fossilis*<br>(n=26, 14.96) | 4.39±0.05<br>=3.41 | 3.96±0.04<br>=3.788 | 5.16±0.04<br>P=2.90 |

(± SE, Figures within parenthesis in the Prey item column indicates the sample size and mean dry wt. in g of the sample. P= Profitability index)

This indicates that Profitability of longer prey like *M. cuchia* is low and it was higher for the group murrels. (Table : 6).

Foraging techniques: a) Visual search and Probing(VSP)-the GAS walks in water with visual search and using its bill (59.87%); b) Active Groping (AG)- the GAS uses its bill submerged while walking slowly (20.21%); c)Pecking and Probing (PP)-the GAS uses its foot to search prey and keeping its bill submerged it walks very slowly (10.36%) and d) Visual search(VS). The GAS walks in water with visual search and thereafter use bill (9.55%) of the total number of fish (n=1256) preyed. The technique d and c are considered as tactile method.

### Assessment of foraging rate and success

The foraging rate, success rate, probe rate and peck rate of GAS varies from wetland to wetland. The study was done by taking five minutes as a bout and later calculated the feeding rate and feeding success per minute. They started walking to the wetland from early morning (approximately 4:30 am during summer), followed by hunting approximately by 5:30 p.m. at the special feeding spot. In the garbage dump, the GAS arrives almost at 4:30 am and start feeding after 1-1.5 h of their arrival. Correlation coefficient matrix (r) analysis showed that PHT:FL= 0.706; PHT:WL=0.860; FS:PK= 0.842; ST: PK=0.843; FS:PR=0.650; FS:ST=0.966 are positively correlated while PR:PK= −0.802; WL:PK= −0.883; WL:FS =−0.839; PR:ST= −0.522; WL:PR= −0.784; WL:PK= −0.883 are negatively correlated. (PHT= Prey handling time in minute; FL= Fish Length in cm; WL=Water level; PK= Peck; ST= Step; PR=Probe; FS=Foraging Success). Feeding rate, success rate in relation to steps, peck and probe has beenpresented in the Table 7 in different habitats used by GAS. The bird deployed tactile method at 79.45%, tactile and visual at 10.98% and visual method at 9.55% in different habitats except garbage dump where it deployed only visual mode of foraging.

**Table 7:** Feeding rate, success rate, steps, peck rate, probe rate (all bird per minute) in Paddy forage field, Swampy wetland and Shallow water during 2012 to 2017 (n=1030; ± SD)

| Types of Habitat | Feeding success /min (capture/min) | Feeding success rate/min | Steps/min | Peck/min | Probe/min |
| --- | --- | --- | --- | --- | --- |
| Paddy field<br>n=350 | 0.68±0.06 | 0.017±0.10 | 25.41±1.44 | 18.23±1.23 | 20.23±2.33 |
| Swampy wetland<br>n=259 | 0.87±0.04 | 0.03±0.05 | 15.23±0.06 | 9.92±0.06 | 17.7±2.06 |
| Shallow Water<br>n=421 | 1.66±0.04 | 0.054±0.06 | 19.24±0.09 | 20.45±0.06 | 10.05±1.23 |
| Total=1030 | 1.03±0.01 | 0.03±0.10 | 22.80±0.03 | 17.33±0.12 | 19.31±3.02 |

Principal Component Analysis: (Fig 3) Component 1 (72.5%) and 2 (26.5%) shows the highest degree of covariance which presents the highest information for the variables. Here the Component 1 Fish Species and the Component 2 is the Site Depth in the area is sharing the maximum covariance with rest of the components as shown in the Screen plot. Component 1 and 2 are correlated in terms of Site 1, 2 and 3 are positively correlated with the fish species found in the area as compared to other prey, *Monopterus cuchia* and *Notopterus chitala* with respect to their larger size and weight. Component 3 is the Prey Size which shows the very less covariance i.e. 0.4% with the rest of the components. The component 3 is the foraging by GAS positively correlate with the prey size greater than 5cm. Component 3 also shows negative foraging correlation with the Prey size greater than 8cm and 15 cm. The Prey full size with beak and above shows negative relation with the Site depth and fish species, but shows good prey for GAS during March to July. Component 4 (0.3%) is the monthly/seasonal foraging activity of the GAS from March to July seems to be same and are less active but from August to December shows the highest foraging activity. The degree of covariance is lesser among the Component 3 and 4 and may be the reason behind it is showing in the negative axis.

Foraging techniques: a) Visual search and Probing(VSP)-the GAS walks in water with visual search and using its bill (59.87%); b) Active Groping (AG)- the GAS uses its bill submerged while walking slowly (20.21%); c)Pecking and Probing (PP)-the GAS uses its foot to search prey and keeping its bill submerged it walks very slowly (10.36%) and d) Visual search(VS)-The GAS walks in water with visual search and thereafter use bill (9.55%) of the total number of fish (n=1256) preyed. The technique d and c are considered as tactile method.

### Assessment of foraging rate and success

The foraging rate, success rate, probe rate and peck rate of GAS varies from wetland to wetland. The study was done by taking five minutes as a bout and later calculated the feeding rate and feeding success per minute. They started walking to the wetland from early morning (approximately 4:30 am during summer), followed by hunting approximately by 5:30 p.m. at the special feeding spot. In the garbage dump, the GAS arrives almost at 4:30 am and start feeding after 1-1.5 h of their arrival. Correlation coefficient matrix (r) analysis showed that PHT:FL= 0.706; PHT:WL=0.860; FS:PK= 0.842; ST: PK=0.843; FS:PR=0.650; FS:ST=0.966 are positively correlated while PR:PK= −0.802; WL:PK= – 0.883; WL:FS =−0.839; PR:ST= −0.522; WL:PR= −0.784; WL:PK= −0.883 are negatively correlated. (PHT= Prey handling time in minute; FL= Fish Length in cm; WL=Water level; PK= Peck; ST= Step; PR=Probe; FS=Foraging Success). Feeding rate, success rate in relation to steps, peck and probe has been been presented in the table 7 in different habitats used by GAS. The bird deployed tactile method at 79.45%, tactile and visual at 10.98% and visual method at 9.55% in different habitats except garbage dump where it deployed only visual mode of foraging.

## Discussion

The foraging ecology of GAS is governed by its morphology, food availability and different food parameters (Mini, 2012) which have primarily been recorded in 6 wetlands and in the lone garbage dump at Boragaon, Guwahati (Table 2). The aerial distance (Table 1) of GAS for foraging ground ranged between 0.9 to 15 km from the breeding colony could well be considered as the effective foraging habitats. The GAS as such has no records in its foraging pattern, yet many workers (Newton, 1986, Alonso *et al*., 1994) had been able to derive certain observations on other wading birds based on the Foraging theory of higher use of sites with greater food availability for their benefit through energy gain (Krebs *et al*., 1983, Stephens and Kreb, 1986). The foraging habitat of wood stork *Mycteria Americana* forages in a habitat with the highest abundance of fishes (Gimeses and Anjos, 2011). Foraging ecology being the most advanced area of stork biology is well studied in wood stork and in black necked stork (Maheswaran and Rahmani, 2002) in terms of seasonality of prey and habitat characteristics. The group ciconiiformes is very well adapted to the monsoon and receding monsoon (Kushlan *et al*.,1985) and sensitive to the seasonal variation in the habitat i.e water level (Gawlik, 2002) as well as prey availability(Del Hoyo *et al*.,1992). Food availability in different habitats was affected by changes in the water level. The Hooded Crane adjusted its foraging patterns and made full use of the three available types of habitat in order to acquire enough food in response to fluctuations in the water level (Zang et al, 2015). Food is a limiting factor during the breeding period in context with the colonial nesting breeders as well as select nesting trees within their feeding territory or feed within their nesting territory and thus minimize the number of foraging trips and increase the vigilance for predators (Butler et al., 1995). Distance of foraging habitats from the breeding site is a significant consideration during the breeding season, when stork must arrive to the breeding site on time to care for their eggs and chicks (Coulter and Bryan, 1993). The distance covered by foraging birds may affect daily energy expenditure of individuals (Baveco et al., 2011), well studied in colonial nesting birds (Bryan and Coulter 1987; Bryan *et al*., 2012). The storks reduces forging travel time in order to save energy to attend their nest, therefore foraging distances are important factors for breeding birds like GAS, and all the foraging sites are within 15.48 km from the breeding zone. It can be assumed from the present study that the Greater Adjutant forages within a range of 15 km from their nesting location, however, transmitter or telemetry study might provide more accurate information on this aspect.

The body morphology is an important factor in limiting the range of foraging behaviour of wading birds (Norazlimi and Ramli, 2015). Body size explains much of the interspecific variation in the physiology, behavior, and morphology of birds, such as metabolic rate, diet selection, intake rate, gut size, and bill size (Mini, 2012). The bill length of GAS (33 cm) and its dagger shape plays a significant role in foraging behaviour, habitat selection and choice of diet (Pierre, 1994) since the longer bill is related with probing and sweeping movement in the wetlands (Friedman *et al*.,2019), while the shorter bills are associated with routing and pecking at the substrate surface (Barbosa and Moreno, 1999).

The foraging technique and PHT of the black-necked stork (*Ephippiorhynchus asiaticus*) showed that the storks foraged adopted a particular technique to procure food. Black-necked storks mostly foraged using a tactile technique (>90%), but sometimes foraged visually. When the water level was estimated to be less than 60 cm, the storks foraged using tactile techniques (Table 3). The shape of bill is either straight or curved influences the foraging techniques (Nebel *et al*., 2005) and the general morphological requirements of wading birds necessitate the bill to be long and narrow, but not very slender and penetration portion should be flattened either vertically or horizontally (Norazlimi and Ramli. 2015). Unlike curved, the GAS has straight, thick and slightly upturned bills forage visually, on fish and a variety of other items including at garbage dumps could be a specialized for digging large eels from the mud (Thomas, 1986, Hancock *et al*. 1992)), adapted for exploiting eels could prove as critical resource for the natural history and survival of the species (Borjas, 2004). In fact, eels may not constitute a relatively large proportion of the diet against around 19 % of the total prey of Jabirus stork, however, Jabirus preferred eels since they are easier to handle than large fish, and perhaps, eels might be ingested entirely without much effort (Borjas, 2004), which could not be established in GAS. It could be argued that dagger shaped bill of GAS helps in tactile foraging in wetlands and also used in other items like carcasses, viscera etc in rubbish dump. The relationship between prey size and handling time (*ht*) was investigated in the Painted Stork by using video techniques (Kalam and Urfi 2008) against the direct observation in this study (Table 6). suggest small prey is easily swallowed, negligible time is spent in handling but with increasing prey size, *ht* increases exponentially. The tibia and tarsus length are correlated with water depth. The GAS foraged in wetlands both in breeding (September to April) and non-breeding seasons (May to Aug) and an adult GAS has adopted to forage up to average of 85 cm of water depth up to the end of tibia (Fig-1)). Sub adult bird had a mean full leg length of 75 cm. Study indicated that the GAS preferred to forage much up to the half of its leg size followed by one third and one sixth of its leg length (Fig 2). Both the adults and sub adult birds were also found to forage up to its tibia as per the availability of the resources.(Table 4, Fig 2). GAS arrives at garbage dump at around 4:30 am and start foraging from 5:30 am but was found to be more active from around 6 am in the garbage dump and became very much adapted to the arrival of garbage trucks. As soon as the garbage trucks arrive, they congregate in the garbage and their feeding frequency increases. The feeding strategy of the Greater Adjutant Stork differs with time during the day in the wetlands. The success rate and peck rate of GAS were positively correlated in different habitats of wetlands. Kushlan (1978) described that the wading birds defend a feeding territory while feeding in groups, although this defended the area may be reduced to each bird’s individual distance, defined by the distance it can reach with its head and neck. Both intra and inter specific competition were seen during foraging. The Greater Adjutant Stork in some observations were found to chase a number of wading birds in garbage dumps and wetlands during foraging. The crucial factor for success is to access foraging habitat (Alonso *et al*., 1994; Olsson and Bolin, 2014) and this has even been confirmed experimentally (Hilgartner *et al*., 2014).

**Fig 1:**
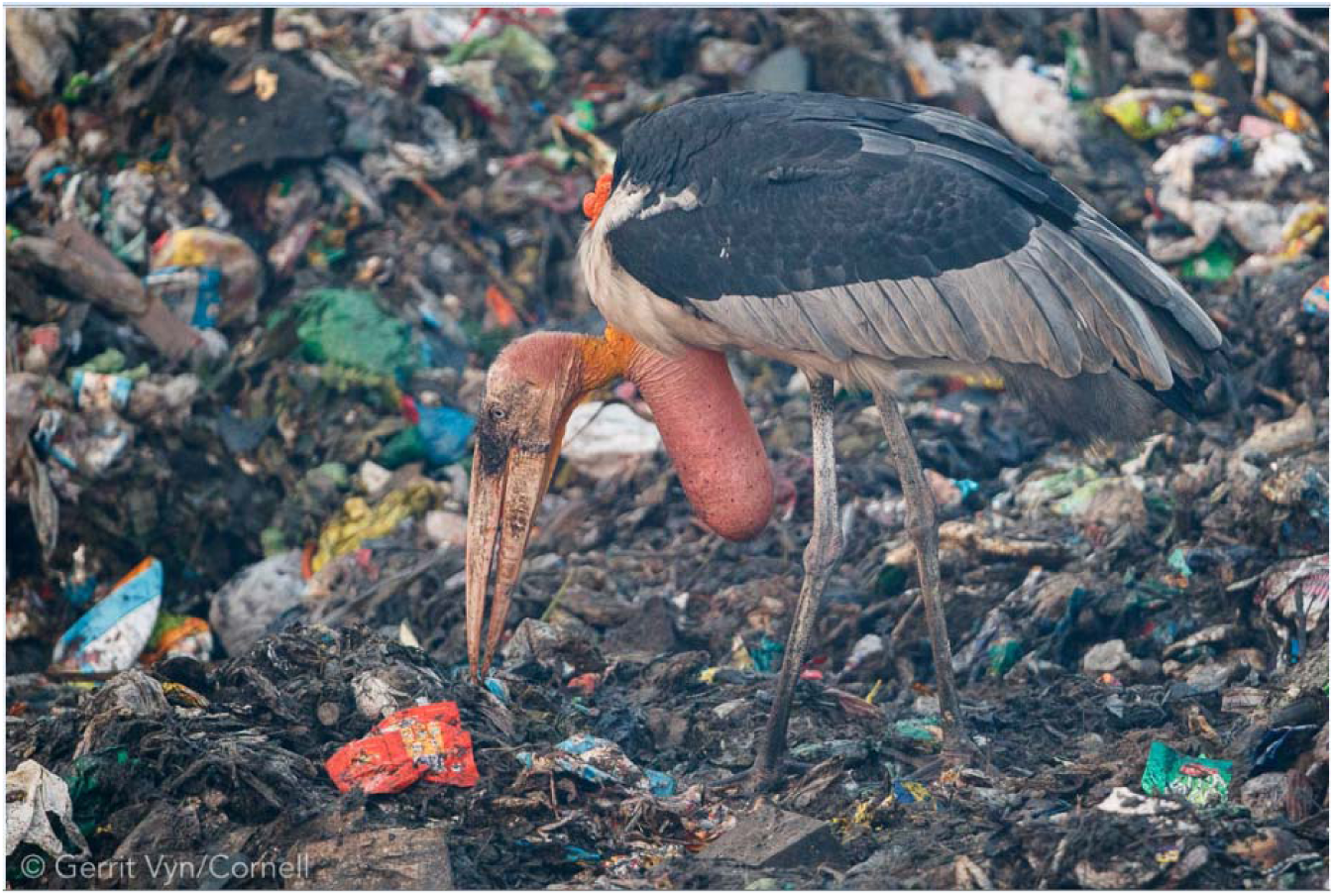
Greater Adjutant forages in a city garbage dump.

**Fig 2:**
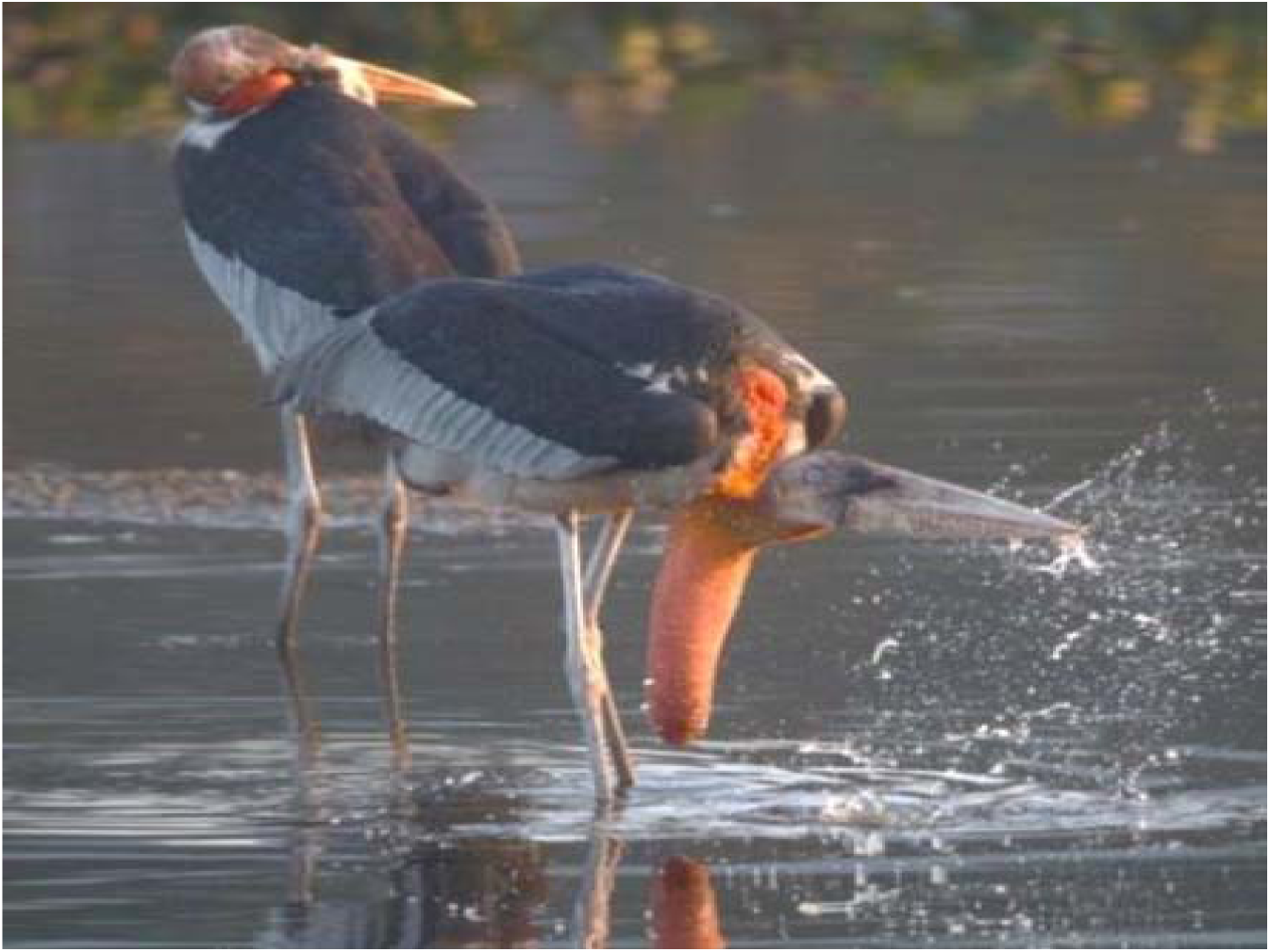
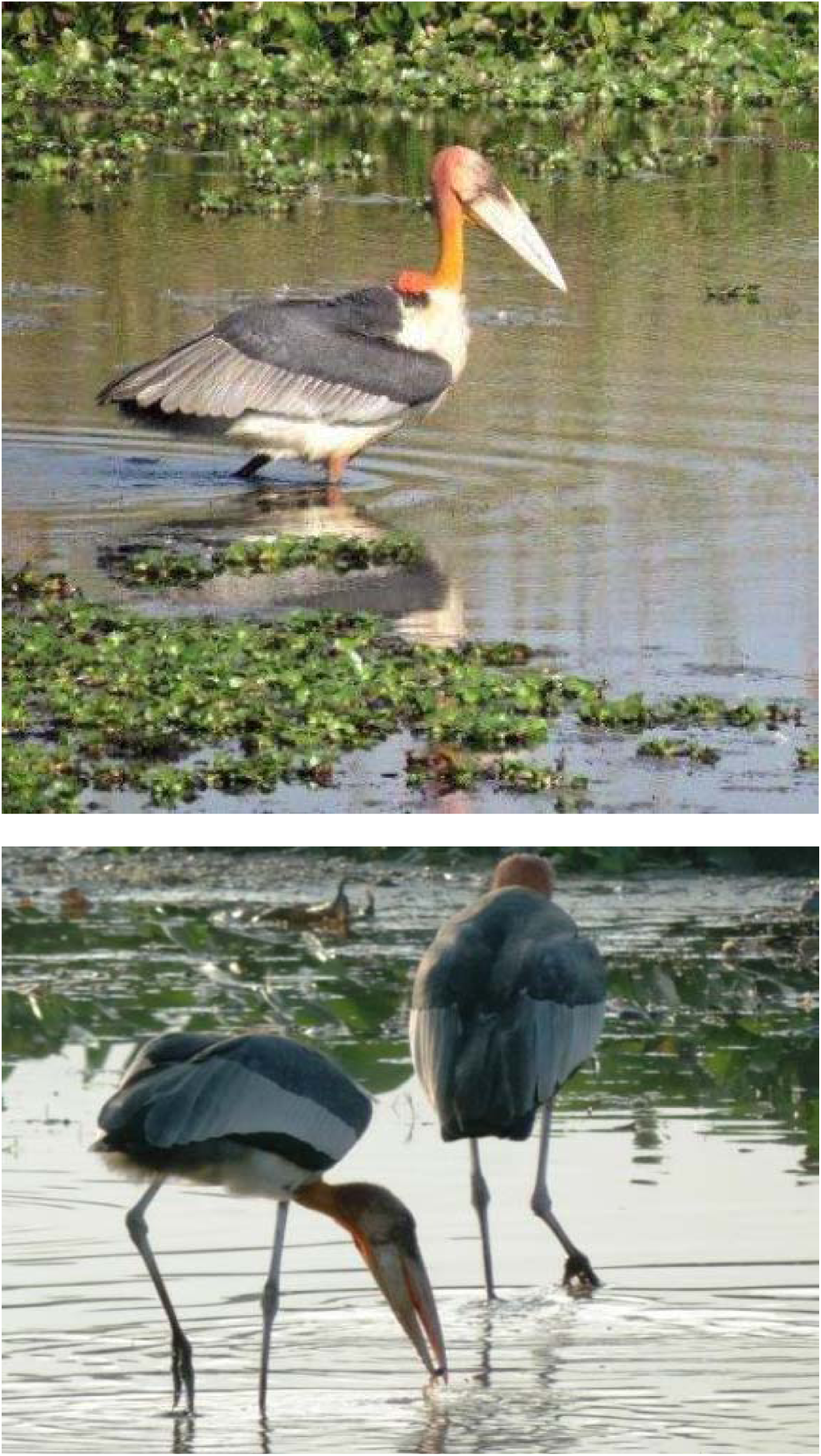
Forages at wetlands coveniently.

**Fig 3:**
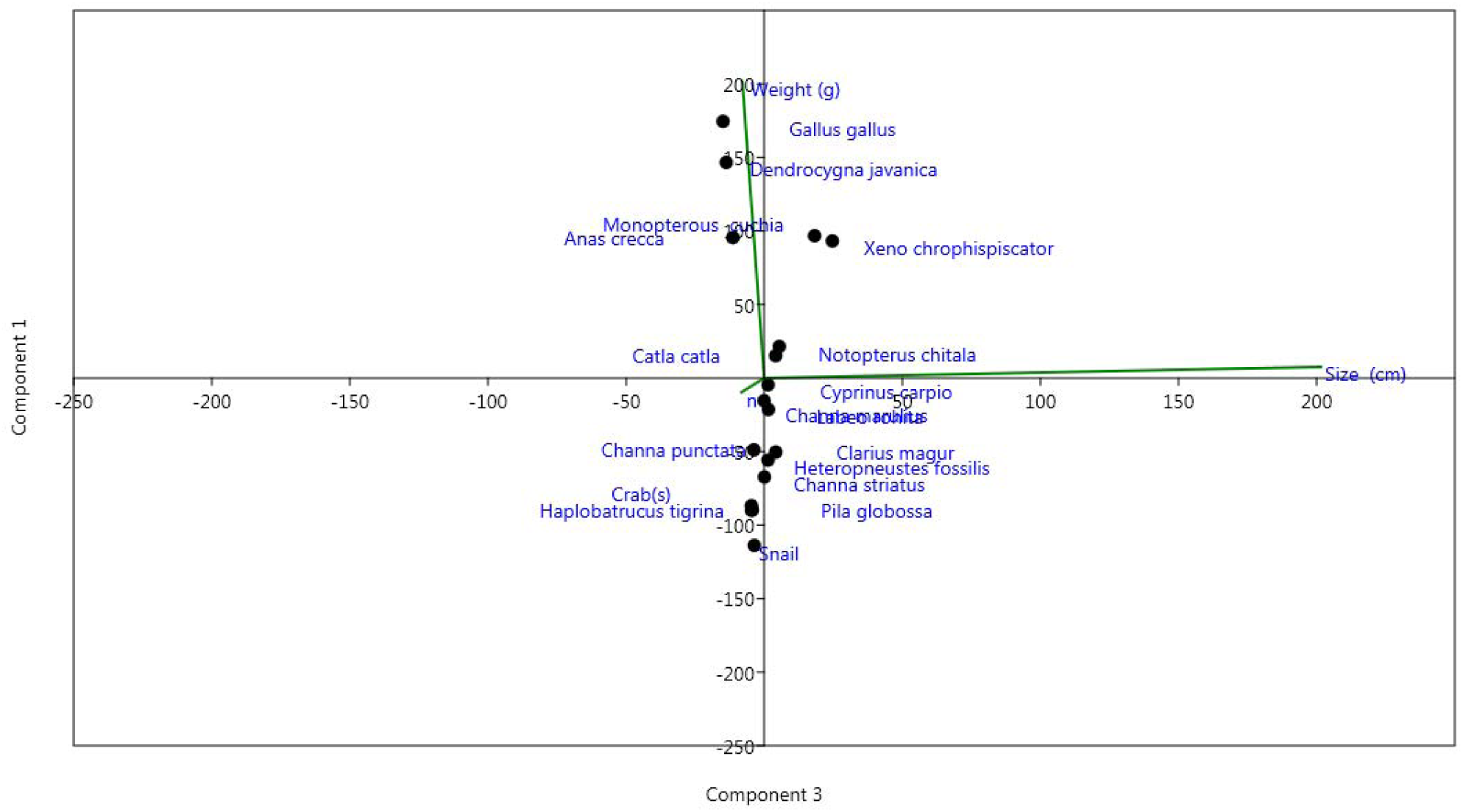
PCA analysis showed the size of the prey and water depth at wet lands in relation to the forages of greater adjutant stork.

Determination of the dietary composition of wild birds is essential for understanding how the GAS interacts with their habitats and consequently for identifying their preferred food types (Zang *et al*.,2017). The diet spectrum of GAS recorded from the regurgitated food analysis showed that Channa and Cyprinids occupies the highest numbers (Table 5). GAS often found to swallow domestic ducks and chickens, Cormorant, Shovellar, Lesser whistling teal, Cotton teal etc. Gut content analysis revealed the presence of unusual materials like nails, hoof, blades etc in the stomach of eight nest fallen died chicks of GAS. This may be due to the constant feeding on the unsegregated garbage dump food. Foraging of White Storks *Ciconia ciconia* on rubbish dump is a common behaviour in northern Africa and the Middle East associated to the wintering of European White Storks in new areas closer to their breedings time in South Africa, where a local breeding population has established (Ciach and Kruszyk, 2010). White Storks *Ciconia ciconia* foraged in a rubbish dump more in non-breeding season than in breeding season due to the constant availability of food resources has facilitated year round nest use (Gilbert et al.,2016) influencing their home ranges and movement behaviour and as such the GAS might be adapted for easier food in garbage dump. The rapid development of garbage dumps may have major consequences for the future ecology of Stork Greater Adjutant.

The foraging technique and prey-handling time of the black-necked stork (*Ephippiorhynchus asiaticus*) showed that the storks foraged adopted a particular technique to procure food. Black-necked storks mostly foraged using a tactile technique (>90%), but sometimes foraged visually. When the water level was estimated to be less than 60 cm, the storks foraged using tactile techniques (Maheswaran and Rahmani, 2008). The GAS being mostly piscivorous are efficient tactile forager similar to that of the painted stork *Mycteria leucocephala* (Kalam and Urfi, 2007). Alonso et al. (1994) suggest that storks followed simple rules of thumb based on flock size, rather than on the more complicated food availability estimations required by central place foraging models. This study in all probability has been able to establish base line information in terms of conservation of this endangered species.

